# TopHap: Rapid inference of key phylogenetic structures from common haplotypes in large genome collections with limited diversity

**DOI:** 10.1101/2021.12.13.472454

**Authors:** Marcos A. Caraballo-Ortiz, Sayaka Miura, Maxwell Sanderford, Tenzin Dolker, Qiqing Tao, Steven Weaver, Sergei L. K. Pond, Sudhir Kumar

## Abstract

**Motivation:** Building reliable phylogenies from very large collections of sequences with a limited number of phylogenetically informative sites is challenging because sequencing errors and recurrent/backward mutations interfere with the phylogenetic signal, confounding true evolutionary relationships. Massive global efforts of sequencing genomes and reconstructing the phylogeny of SARS-CoV-2 strains exemplify these difficulties since there are only hundreds of phylogenetically informative sites and millions of genomes. For such datasets, we set out to develop a method for building the phylogenetic tree of genomic haplotypes consisting of positions harboring common variants to improve the signal-to-noise ratio for more accurate phylogenetic inference of resolvable phylogenetic features.

**Results:** We present the *TopHap* approach that determines spatiotemporally common haplotypes of common variants and builds their phylogeny at a fraction of the computational time of traditional methods. To assess topological robustness, we develop a bootstrap resampling strategy that resamples genomes spatiotemporally. The application of *TopHap* to build a phylogeny of 68,057 genomes (68KG) produced an evolutionary tree of major SARS-CoV-2 haplotypes. This phylogeny is concordant with the mutation tree inferred using the co-occurrence pattern of mutations and recovers key phylogenetic relationships from more traditional analyses. We also evaluated alternative roots of the SARS-CoV-2 phylogeny and found that the earliest sampled genomes in 2019 likely evolved by four mutations of the most recent common ancestor of all SARS-CoV-2 genomes. An application of *TopHap* to more than 1 million genomes reconstructed the most comprehensive evolutionary relationships of major variants, which confirmed the 68KG phylogeny and provided evolutionary origins of major variants of concern.

**Availability:** *TopHap* is available on the web at https://github.com/SayakaMiura/TopHap.

## 1 Introduction

The global health emergency caused by the SARS-CoV-2 coronavirus has catalyzed an unprecedented effort to sequence millions of genomes from all around the world and their analysis to reveal its origins and evolution (Andersen *et al*., 2020; Kumar *et al*., 2021; Rambaut *et al*., 2020). However, applying classical phylogenetic methods to infer the global SARS-CoV-2 phylogeny has been challenging (Kumar *et al*., 2021; Morel *et al*., 2020). This is partly because phylogenetically informative sites are relatively rare due to a low mutation rate and a short evolutionary period of the outbreak. Genome sequences contain random and systematic sequencing errors, which compete with informative phylogenetic variation and mislead phylogenetic inference (Kumar *et al*., 2021; Morel *et al*., 2020; Pipes *et al*., 2020). Consequently, the application of standard phylogenetic methods to the multiple sequencing alignment (MSA) of SARS-CoV-2 genomes has produced many equally plausible phylogenies, particularly when reconstructing early mutational history and the root of the SARS-CoV-2 phylogeny (Nie *et al*., 2020; Pipes *et al*., 2020; van Dorp *et al*., 2020).

Kumar *et al*. (2021) reconstructed a mutation tree using shared cooccurrence patterns of mutations occurring in >1% of isolates. They applied and advanced a maximum likelihood method (SCITE, Jahn *et al*., 2016) that models false-positive and false-negative variant detections while assuming that recombination is absent (Jahn *et al*., 2016; Kumar *et al*., 2021). They reported success deciphering the earliest phases of SARS-CoV-2 evolution, including the most recent common ancestor (MRCA) genome, using common variants observed in the early stages of SARS-CoV-2 evolution. Based on the success in building the mutation tree using common variants, we hypothesized that it should be possible to build a reliable molecular phylogeny of major SARS-CoV-2 haplotypes by filtering out all genomic positions at which no minor allele rose to a frequency greater than 1%. Such filtering should effectively reduce the effect of the noise in making molecular phylogenetic inferences using standard approaches (e.g., the maximum likelihood [ML] method). If successful, one would prefer a traditional phylogenetic approach because it can better handle multiple substitutions at the same site (homoplasy) and use outgroup sequences more effectively than the mutation tree approaches.

However, excluding alignment sites with only low-frequency variants followed by applying a standard phylogenetic approach on remaining sites was unsuccessful. An example illustrates the problem in **figure 1**. The ancestral genome contains only three polymorphic positions where derived alleles occur at high frequencies (#1, #2, and #3; **Fig. 1a**). In this case, we expect to see at most four correct haplotypes in the absence of noise: three mutant strains (H1, H2, and H3) and one ancestral haplotype. The addition of a small number of sequencing errors and homoplasy generate additional haplotypes (e.g., H4, H5, and H6) that occur with very low frequency but still mislead an ML analysis (**Fig. 1b**). ML phylogenies place two spurious haplotypes (H5 and H6) near the root of the tree (**Fig. 1c**), albeit without significant statistical support.

**Figure 1.**
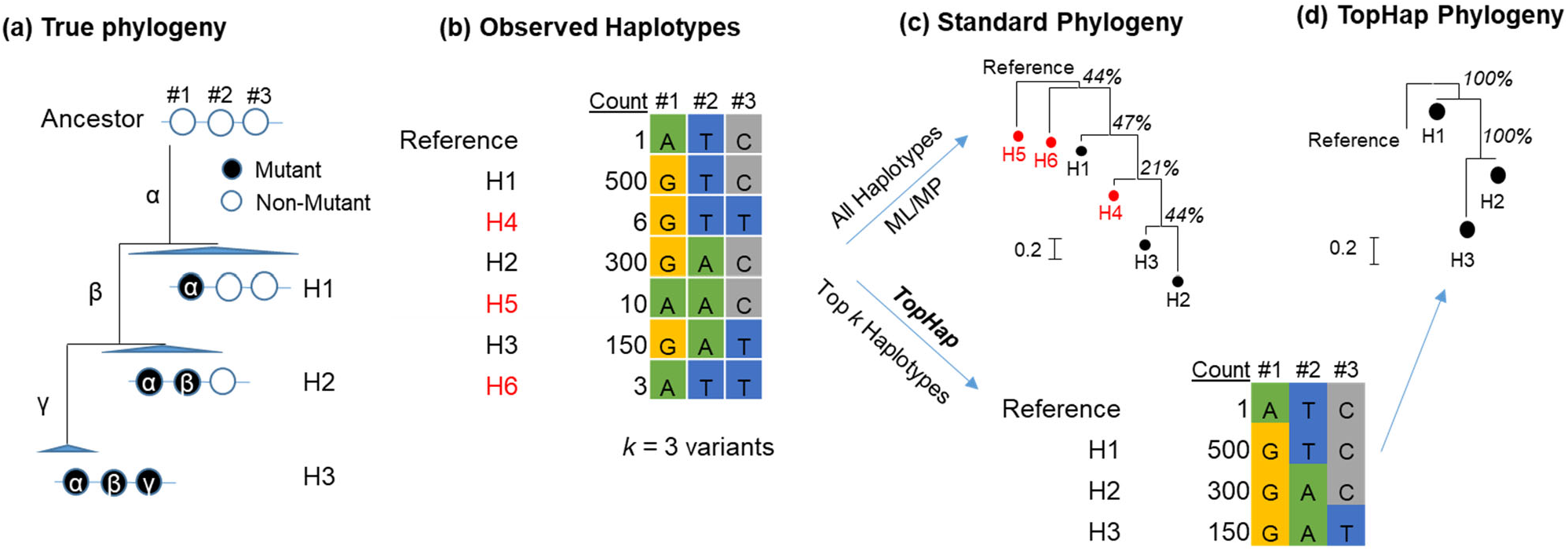
Traditional phylogenetic approach versus the new *TopHap* approach to manage a dataset that contains many sequences with few variants. (**a**) The true tree showing three simulated haplotypes. In this example, three mutations (α, β, and γ) occurred sequentially and gave rise to haplotypes, H1, H2, and H3. The size of triangles at each tip is proportional to the number of genomes containing these haplotypes. (**b**) Phylogenetic approaches use a multiple sequence alignment, simplified here with only three informative variants. Due to sequencing errors, a few spurious haplotypes would be observed (red letters, i.e., H4–H6) in low haplotype frequencies (0.3%–1%). Inclusion of these spurious haplotypes misguides the maximum likelihood (ML) and other methods and distorts the evolutionary inference. (**c**) Results based on a typical ML approach would suggest that the spurious haplotypes H6 and H5, were the first to arise. The bootstrap support for all the branching patterns is low (21%–47%) because each branch is only one mutation long, a situation where the bootstrap method is known to be powerless (see text). (**d**) The *TopHap* approach was able to infer the correct tree because it uses high-frequency haplotypes to infer a phylogeny; the haplotypes with low frequencies – which are likely spurious due to sequencing errors – are removed.

However, this behavior is rectified when one removes rare haplotypes (**Fig. 1d**). This observation prompted us to develop a simple filtering procedure to identify *common (top) haplotypes* of *common variants* for molecular phylogenetic analysis. We first present this filtering process (the *TopHap* approach) and then apply it to infer the early evolutionary history of SARS-CoV-2 by using 68,057 genomes previously analyzed by Kumar *et al*. (2021) for a direct comparison of the *TopHap* phylogeny with the mutation tree generated by using patterns of co-occurrence of variants.

## 2 Methods

### 2.1 The *TopHap* approach

As input, *TopHap* uses an MSA of genomes (*n* genomes and *m* base positions; **Fig. 2a**). The first step is the selection of common variants by specifying a desired minor allele frequency threshold (e.g., *maf* > 1%) without using any reference genomes. All alignment sites containing at least one allele with a frequency greater than *maf* and another allele with a frequency less than *1-maf* are marked for inclusion (*k* variant positions). Every genome is then reduced to a haplotype on *k* retained sites. Next, unique haplotype sequences are identified, and their frequencies tallied.

**Figure 2.**
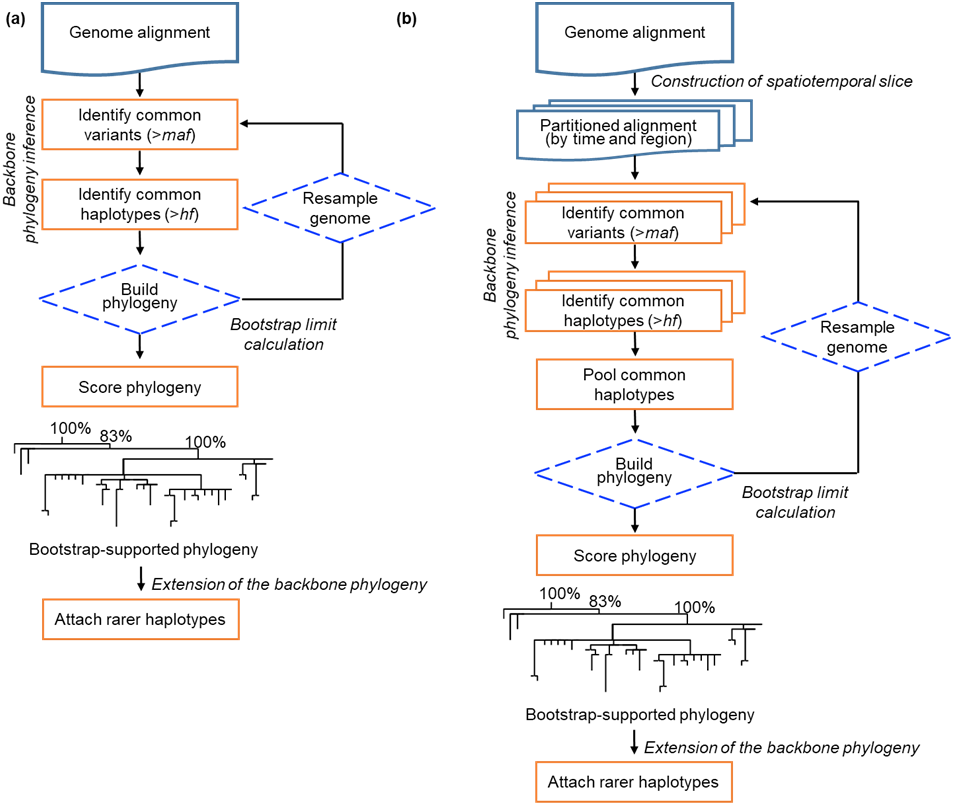
Overview of the *TopHap* approach. Input to *TopHap* is an alignment of genome sequences (*n* sequences, *m* bases each). *TopHap* flowchart (**a**) without any spatiotemporal sampling information and (**b**) with consideration of spatiotemporal information are shown. In the *TopHap* algorithm, we first identify high-frequency variants (>*maf*) and produce a restricted alignment with *n* sequences and *k* bases. Next, we identify high-frequency haplotypes (>*hf*) and produce a new alignment of *h* haplotypes, each with *k* bases. These haplotypes are subjected to phylogenetic inference. To compute bootstrap confidence limits, we resample *n* haplotypes with replacement to form a replicate *n*×*k* dataset, identify high-frequency haplotypes (>*hf*) and then infer a phylogeny. This process is repeated 100 times and a bootstrap consensus phylogeny is inferred. When the analyses are conducted with the consideration of spatiotemporal (or other type) annotation to construct subsets (**panel b**), variants and haplotypes are identified for each slice separately and then they are merged prior to phylogenetic analysis. For the SARS-CoV-2 68KG dataset, we classified sequences based on their sampling months and sampling countries.

*TopHap* selects the top *h* haplotypes given a desired *hf* frequency cutoff. Now, the MSA contains *h* haplotypes, each *k* base positions long and tagged with its frequency. Outgroup genomes are added into the MSA by converting them into haplotypes containing only *k* selected positions. This reduced MSA is subjected to the phylogenetic analysis using standard methods, e.g., Maximum Parsimony (MP) or ML methods with models for analyzing datasets containing only polymorphic positions (Lewis, 2001; Schrempf *et al*., 2016; Stamatakis, 2014). This will produce the *TopHap* phylogeny of common haplotypes on common variants.

When information on sampling location and time of haplotypes is available, *TopHap* selects variants and haplotypes for each spatiotemporal slice of the dataset that is regionally (e.g., continent, country, or city) and/or temporally (e.g., monthly) partitioned (**Fig. 2b**). Then, all the variants and haplotypes selected in spatiotemporal slices are pooled together. In this case, the same *maf* and *hf* thresholds are applied to each spatiotemporal slice.

#### Calculation of bootstrap support

In the *TopHap* approach, bootstrap branch support for the inferred phylogeny of common haplotypes is calculated by resampling genomes to build bootstrap replicate datasets. This procedure assesses the robustness of the inferred phylogeny to the inclusion/exclusion of haplotypes likely created by sequencing errors and convergent changes that are expected to have relatively low frequencies spatiotemporally because the bootstrap resampling procedure is applied separately to each spatiotemporal slice and the final set of variants and haplotypes are pooled together. This genome resampling approach is different from Felsenstein’s bootstrap approach of resampling sites to build bootstrap replicate datasets, which is not useful for phylogenies in which many branches are expected to have fewer than three mutations. At least three mutations per branch are required to achieve a 95% confidence level even without any homoplasy in the standard bootstrap approach (Felsenstein, 1985; Soltis and Soltis, 2003).

Every bootstrap replicate dataset is subjected to phylogeny reconstruction to generate bootstrap replicate phylogenies. Haplotypes that do not appear in all the bootstrap replicates are then pruned from all the phylogenies. (Of course, one may choose to retain all haplotypes that occur in most of the bootstrap replicates.) Then a bootstrap consensus tree is generated. This consensus tree contains bootstrap support values on the inferred phylogeny.

#### Placement of additional haplotypes into the phylogeny

To place a new genome into the *TopHap* phylogeny, the first step is to transform it into a haplotype of *k* positions used to build the *TopHap* phylogeny. One may use an MP (e.g., UShER (Turakhia *et al*., 2021) and pplacer (Matsen *et al*.,2010)) or ML approach (RAxML-EPA (Berger *et al*., 2011)). When the intent is to place a genome with variant(s) in the genomic position that was not used to build the *TopHap* phylogeny, we rebuild the phylogeny by requiring that the position(s) of interest be always included during the *TopHap* analysis. This process takes only a few additional minutes and produces a phylogeny with haplotypes that contain the variant(s) of interest.

#### Annotations using Nextstrain and PANGO classifications

To compare *TopHap* phylogeny with the Nextstrain classification, we annotated all the *TopHap* haplotypes using the presence and absence of diagnostic Nextstrain mutations (https://Nextstrain.org/ncov). We also assigned a PANGO lineage to each genome in the data using the Phylogenetic Assignment of Named Global Outbreak Lineages (PANGOLIN) software (Rambaut *et al*., 2020). *TopHap* haplotype ID was also assigned for each genome when an observed haplotype was perfectly identical to a *TopHap* haplotype. Here, the same *TopHap* haplotypes may belong to more than one different PANGO lineage. In this case, we paired a *TopHap* haplotype with the major PANGO lineage.

### 2.2 Genome Data Acquisition and Assembly

We obtained an MSA containing 68,057 genomes (hereafter, 68KG) of the SARS-CoV-2 coronavirus from human hosts analyzed in Kumar *et al*., (2021). These genomes were obtained from the GISAID database (https://www.gisaid.org) and covered the period from December 24, 2019, until October 12, 2020. The 68KG alignment was generated after filtering 133,741 SARS-CoV-2 genomes, such that genomes shorter than 28,000 bases and those with many ambiguous bases were removed. Three outgroup coronavirus genomes were added to the alignment: *Rhinolophus affinis* (RaTG13) and *R. malayanus* (RmYN02) bats, and the *Manis javanica* pangolin (MT040335) (Liu *et al*., 2020; Zhou *et al*., 2020). Following the above procedure, we also assembled a dataset containing 1,106,862 genomes (hereafter 1MG) from the GISAID database covering the period from December 24, 2019, to September 11, 2021.

## 3 Results

We stratified sequence isolates by (month of sampling, country) attributes to select variants and haplotypes in the *TopHap* analysis of the 68KG dataset. We used spatial and regional *maf* and *hf* cut-offs of 5% to avoid including problematic variants and haplotypes created by recurrent/backward mutations and sequencing error, particularly because of the small number of genomes available for some spatiotemporal slices. We retained a data stratum that contained at least 500 genomes. When the number of genomes sampled from a country was less than 500, we manually pooled them with adjacent countries with sparse sampling, i.e., all genomes from these countries were pooled when they were located in the same continent. Also, the numbers of genomes in December 2019 and October 2020 were <500, so we pooled them with January 2020 and September 2020 time slices, respectively.

We used the Maximum Parsimony approach to infer *TopHap* phylogeny because the dataset contained only variable sites. One hundred bootstrap replicate analyses were carried out following the procedure outlined in the methods section. The whole *TopHap* analysis was completed in less than an hour, providing the spatiotemporal slices of 68K haplotypes composed of positions with *maf* > 5%. The final tree consisted of 83 variable sites and 39 unique haplotypes (**Fig. 3**). All but three groupings in this tree were highly supported (bootstrap confidence levels > 95%), i.e., many inferences were robust to genome sampling (100 replicates). The other three groupings received >80% bootstrap support. In the *TopHap* phylogeny, many branches were longer than one mutation, and many zerolength branches created multifurcations.

**Figure 3.**
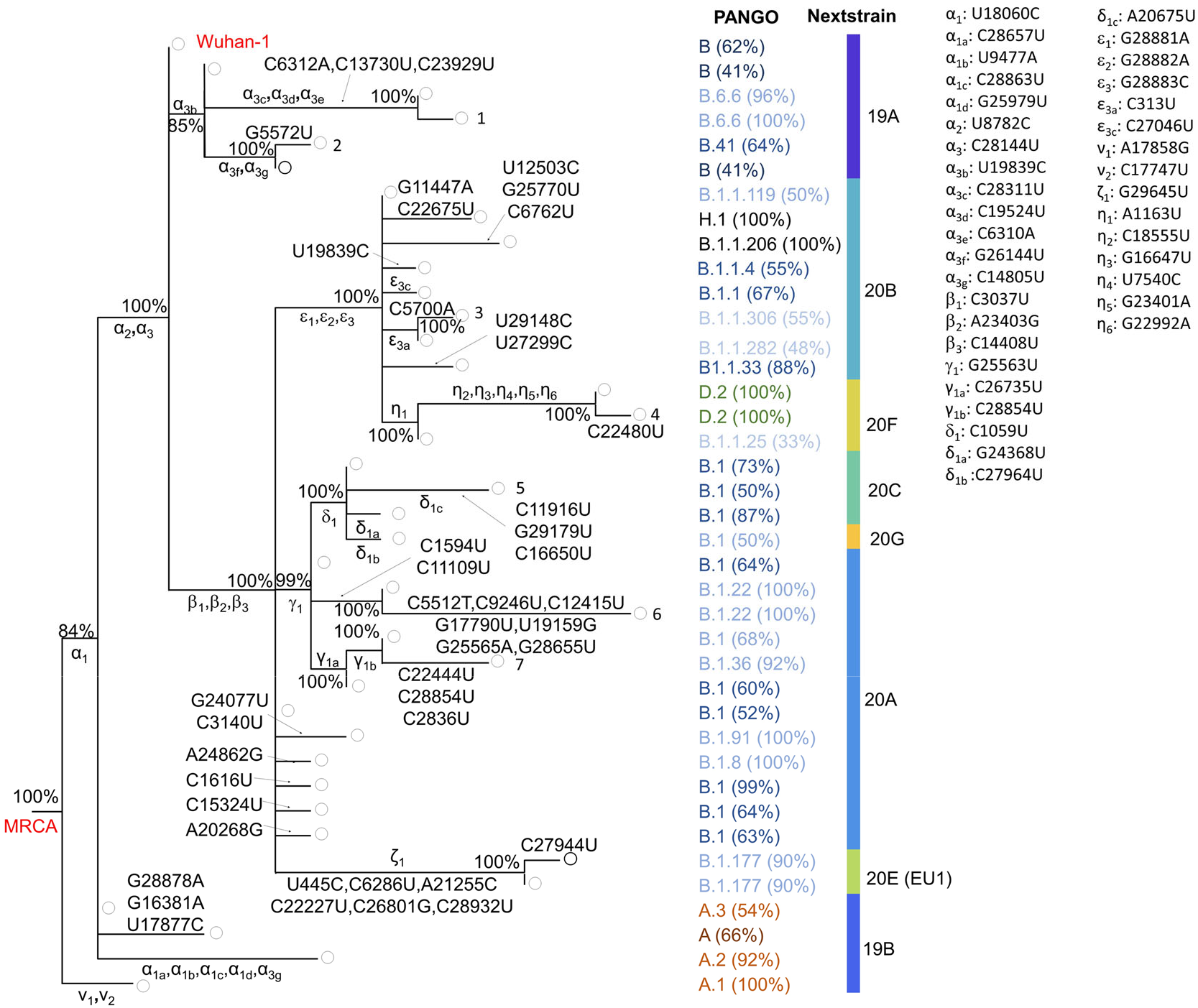
The *TopHap* Phylogeny of 68KG SARS-CoV-2 major haplotypes. (**a**) Numbers near nodes are bootstrap confidence limits derived from bootstrap resampling of genomes. Mutations mapped are shown on branches. When the same mutations were included in Kumar *et al*., (2021), their mutation IDs (Greek symbols) were shown. Their mutations and genomic positions are given in the right side. The Nextstrain clade ID was annotated based on their diagnostic mutations and is provided at the far right. PANGO lineage was annotated for each genome using PANGOLIN software (Rambaut *et al*., 2020). We also annotated *TopHap* haplotype for each genome by comparing its haplotype with *TopHap* haplotypes. When an observed haplotype did not perfectly match any of the *TopHap* haplotypes, we did not assign any for the genome. Using these genome annotations, we paired each *TopHap* haplotype with the major PANGO lineage, and the percentage of genomes containing it is presented in the parenthesis.

### Temporal trends in variant frequencies

The *TopHap* approach in the MP analysis of selected haplotypes does not use the temporal information of genomes (isolation day) nor observed variant frequency. Therefore, a *TopHap* phylogeny allows us to test the concordance between mutation occurrence time and variant frequency and those expected from the phylogeny. We first mapped mutations to every branch in the SARS-CoV-2 phylogeny by reconstructing the most parsimonious ancestral states and minimum evolution. All mutations were mapped unambiguously (**Fig. 3**). In the *TopHap* phylogeny of haplotypes from the early pandemic (up to October 2020), the frequencies of variants mapped onto branches generally decreased from the root to tip on evolutionary lineages (e.g., **Fig 4a**). For example, the mutant bases mapping to the earliest diverging branches in the *TopHap* phylogeny occurred with the highest frequency in the 68KG dataset. Also, the timing of the first sampling date of variants increased on lineages from root to tip (**Fig 4b**). This good concordance with those expected from the inferred *TopHap* phylogeny indicates that the observed patterns are consistent with the clonal evolution of SARS-CoV-2 during the early stage of the pandemic.

**Figure 4.**
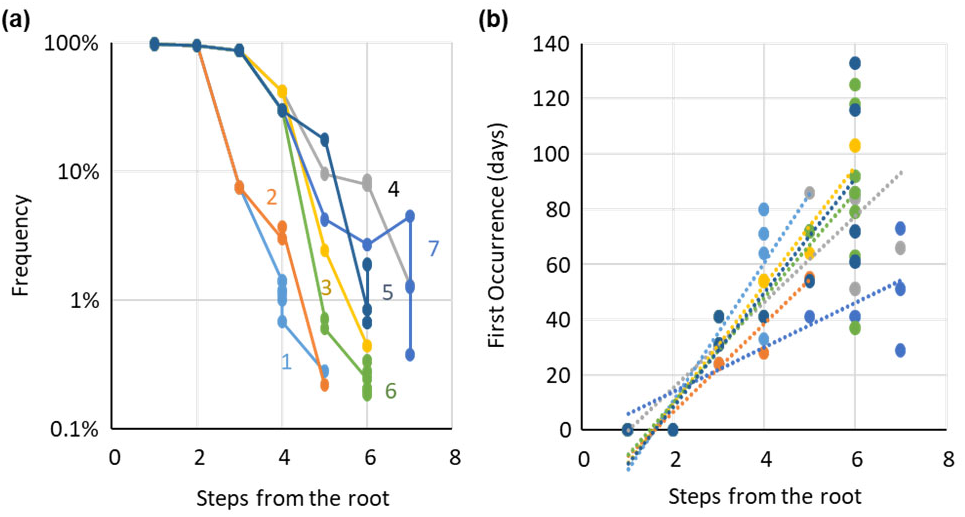
Number of branches from the root to a tip and global mutant nucleotide frequency (**b**) and the first time the mutation was observed (**c**). Selected tip IDs are found in **Figure 3**. The day was counted from the first sample date (Dec 24, 2019).

### Comparing the TopHap phylogeny with the mutation tree

We compared the order of mutations in the *TopHap* phylogeny with the mutation tree inferred using the co-occurring mutations pattern by Kumar *et al*. (2021). This is different from the MP analysis of *TopHap* haplotypes as it treated every variant position independently. For notational consistency, we used Greek symbols for variants when they were the same as those analyzed in Kumar *et al*. (2021, Supplementary Table 2). In total, 49 mutations were shared between 68K *TopHap* phylogeny (83 mutations) and the mutation tree in Kumar *et al*. (2021) (84 mutations). Thirty-four variants were unique to the *TopHap* phylogeny because those variants and corresponding haplotypes rose to high enough frequency (5% frequency) in at least one spatiotemporal slice (locale or month), but their global frequency was less than the 1% required by Kumar *et al*., (2021) in building the mutation tree. Thirty-five mutations analyzed in Kumar *et al*. (2021) were missing from the *TopHap* analysis because they never rose to 5% frequency in any time slice. So, we based our comparison of *TopHap* and mutation tree on 49 variants found in both analyses.

The inferred mutation order from the *TopHap* phylogeny is consistent with the mutation tree, except that some mutations on branches with multiple mutations were not resolved. For example, the mutation tree suggested that α_2_ mutation preceded α_3_, but *TopHap* phylogeny does not resolve this transition because the corresponding intermediate haplotype did not occur with more than 5% frequency in any spatiotemporal slice. This *TopHap* result is reasonable because sequencing error may have created spurious transitional haplotypes (Pekar *et al*., 2021). At the same time, some multi-mutation branches in the *TopHap* phylogeny also correspond well with the unresolved branching order of mutations in Kumar *et al*. (2021), which was suggested to be due to evolutionary bursts (e.g., three β mutations and three ε mutations).

The high concordance of *TopHap* phylogeny inferred, without making any assumptions about the lack of recombination, with the mutation tree inferred using variant co-occurrence patterns suggests that the early phases of SARS-CoV-2 evolution did not involve significant numbers of recombination and co-infections (see also Varabyou *et al*., 2021), which could have, otherwise, resulted in differences between the *TopHap* phylogeny and the mutation tree.

### TopHap analysis with >1 million SARS-CoV-2 genomes

Next, we analyzed a recent snapshot of SARS-CoV-2 genome collection acquired one year after the 68KG data set. After filtering incomplete genome sequences, we obtained an alignment of 1,106,862 genomes to build the 1MG dataset that was 16 times bigger than the 68KG dataset. Nevertheless, the *TopHap* analysis, including 100 bootstrap replicates, was completed in less than 3 hours. Using TopHap with *hf* > 5% and *maf* > 5%, we obtained a reduced MSA of 150 haplotypes with 675 variable sites. Therefore, the number of haplotypes increased only four-fold, and the number of variable sites increased by eight times. The inferred phylogeny from the 1MG data set (**Figure 6a**) was supported with relatively high bootstrap values (>80%) and concordant with the 68KG *TopHap* phylogeny. The order of early mutations (α_1−_α_3_, β_1−_β_3_, ε_1−_ε_3_, γ_1_, δ_1_ and ν_1−_ν_2_) was the same for 1MG and 68KG *TopHap* phylogenies. Therefore, inferences about the early history reported for the 68KG data set were robust to the expanded sampling of genomes.

The 1MG *TopHap* phylogeny contains several branches with many mutations, which often lead to haplotypes that have been designated variants of concern (VOC) by WHO. This includes WHO-ALPHA, WHO-BETA, WHO-DELTA, WHO-ETA, WHO-GAMMA, and WHO-LAMBDA variants. We used the WHO-prefix to avoid conflicting between Kumar et al. notations for mutations and the WHO’s notation for multi-mutation strains. Notably, Kumar et al. mutation identifiers were proposed earlier, so we have retained them. The placement of these VOCs in *TopHap* is like that in the Nextstrain (**Fig. 6b** and **6c**). For example, both Nextstrain and *TopHap* reconstruct WHO-ALPHA, WHO-GAMMA, and WHO-LAMBDA to be sister lineages. Also, the N501Y Spike recurrent mutation (A23063T) occurred independently in WHO-ALPHA, WHO-GAMMA, and WHO-BETA lineages, inferred correctly by *TopHap* (**Fig. 6a**).

### Rooting the tree of SARS-CoV-2 genomes

We find that Nextstrain and PANGO phylogeny broadly agree with 68KG and 1MG *TopHap* phylogenies, except for the root position (**Fig. 3, 5,** and **6**). For example, clade 19A is at the root of the Nextstrain phylogeny, but *TopHap* phylogenies using the bat/pangolin outgroups suggest that Clade 19A is a derived clade. Based on 100 genome resamples, the bootstrap support was modest (>66%) for the root of the *TopHap* phylogeny. But, no bootstrap replicates supported the Nextstrain rooting, and <34% supported the PANGO rooting.

**Figure 5.**
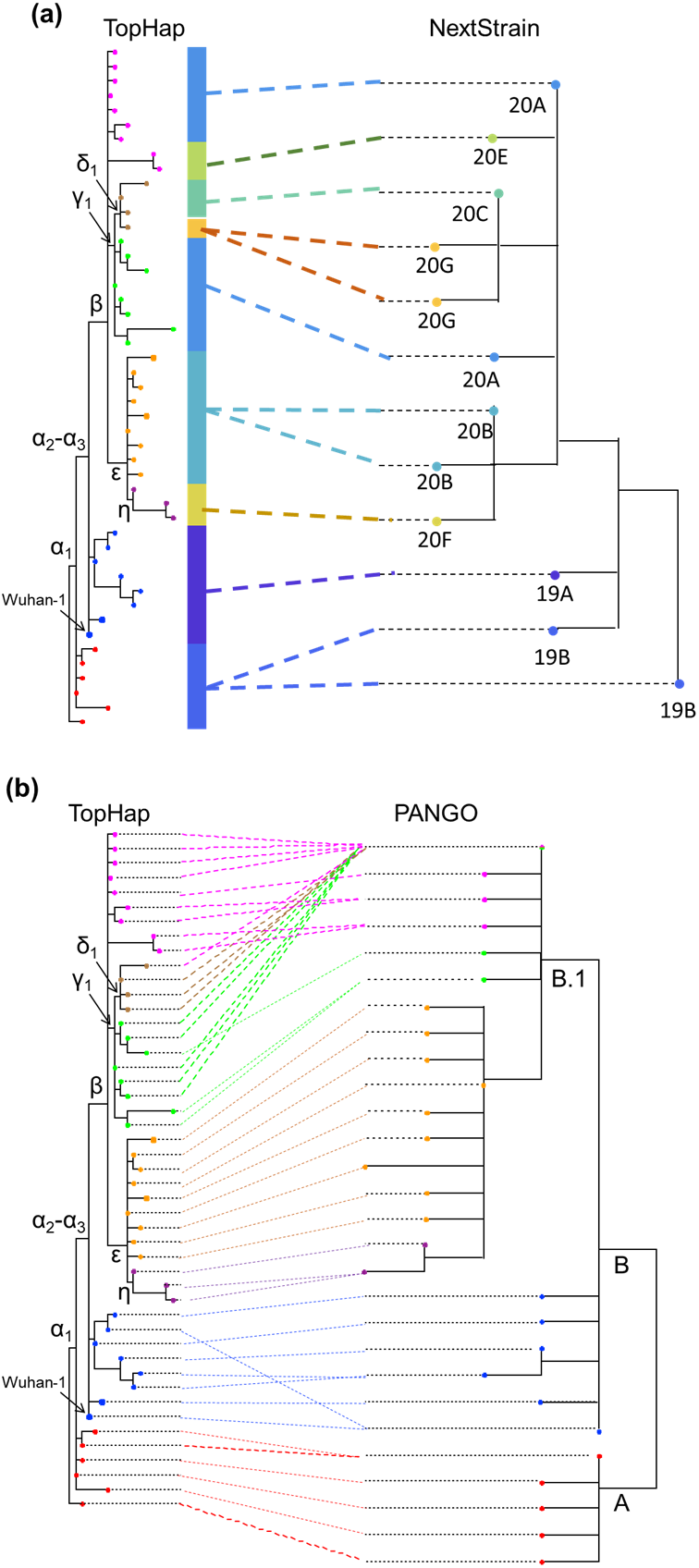
Comparison of TopHap phylogeny with the Nextstrain (a) and PANGO (b) phylogenies. (**a**) Only clades included in the 68KG data are shown. (b) Only PANGO lineages that were included in the TopHap phylogeny were used. Corresponding PANGO IDs are found in **Figure 3.**

**Figure 6.**
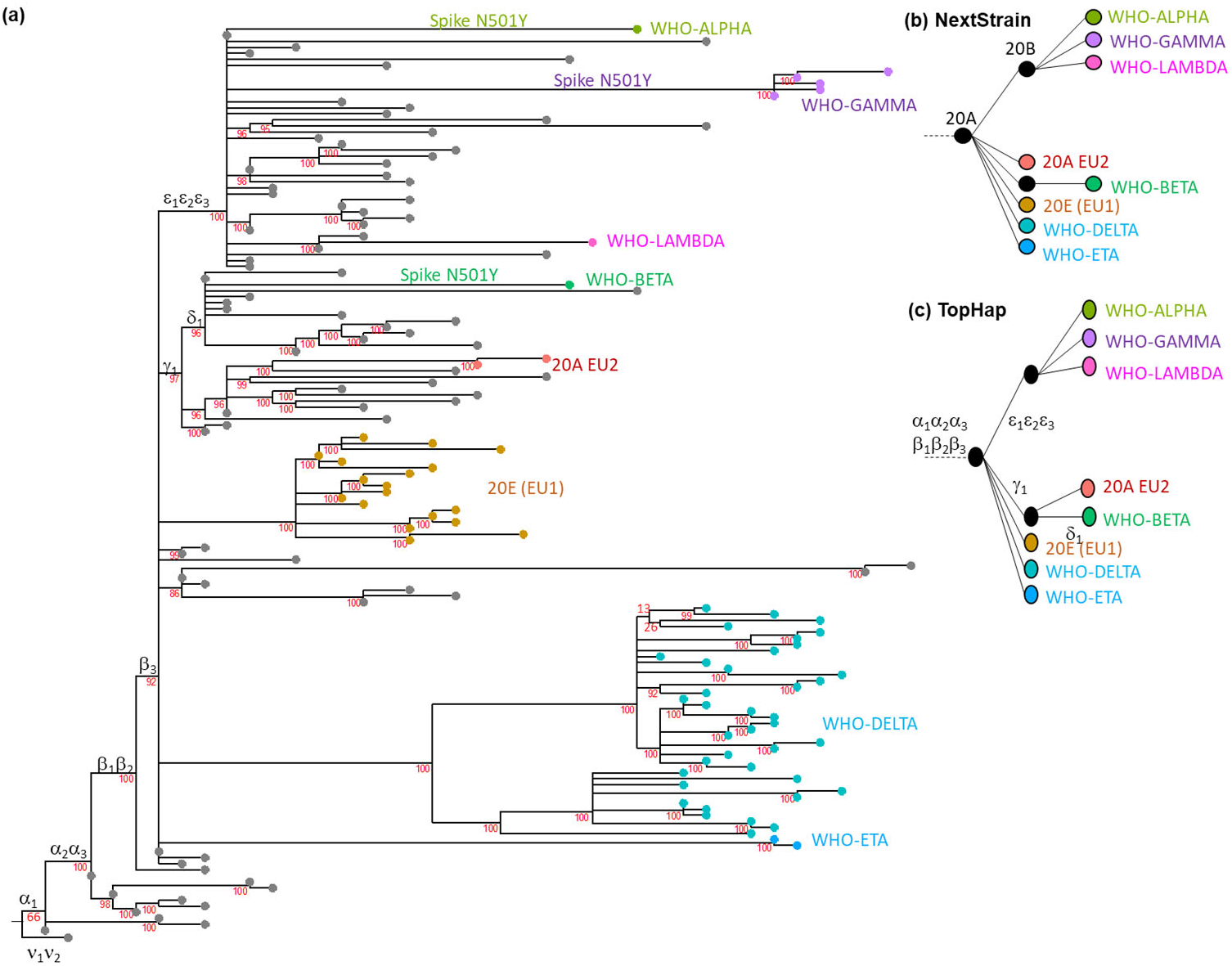
The 1MG *TopHap* Phylogeny. (**a**) Red numbers near nodes are bootstrap confidence limits derived from bootstrap resampling of genomes. Early mutations that were predicted in Kumar *et al*. (2021) are mapped are shown on branches using their mutation IDs (Greek symbols). Their mutations and genomic positions are given in **Figure 3**. The haplotypes with concerning mutations are indicated by using WHO IDs, and 20A EU2 and 20E (EU1) are Nextstrain clade IDs. These haplotypes were identified by annotating PANGO and Nextstrain lineage for each genome. We also annotated *TopHap* haplotype for each genome by comparing its haplotype with *TopHap* haplotypes. When an observed haplotype did not perfectly match any of the *TopHap* haplotypes, we did not assign any for the genome. Using these genome annotations, we paired each *TopHap* haplotype with the major PANGO and Nextstrain lineage, which contained the WHO annotation. We assigned WHO ID when at least one of the annotations indicated it. Evolutionary relationship of lineages with concerning mutations by (**b**) Nextstrain and (**c**) *TopHap*.

The *TopHap* rooting is the same as that implied by the mutation tree reported in Kumar *et al*. (2021). *TopHap* root is also consistent with one of the two preferred roots in Bloom (2021), who analyzed 13 additional partial genomes from only China until the end of January 2020. Key early mutations analyzed in Bloom (2021) contained an additional variable site (genomic position 29095), where the minor base occurred with a too low frequency to be included in the *TopHap* analysis (0.4% in the 68KG). So, we refer to it as mutation *x* (= 29095, U is minor, and C is major) and include it in the 68KG MSA. We also searched for other rare haplotypes to see if any other cluster at or near the root position in the 68KG *TopHap* phylogeny.

We found 936 additional unique haplotypes that occurred in the 68KG dataset more than once. We tested their placement one by one in the *TopHap* phylogeny by using RAxML-EPA (Berger *et al*., 2011). Only two were attached at or near the root. One of them had the same haplotype sequence as that of MRCA and was present in 17 isolates. This haplotype is the proCoV2 sequence reported by Kumar *et al*. (2021), circulating in early 2020. The other haplotype differed from the proCoV2 sequence in two genomic positions (29095 [location of *x* variant] and 18060 [location of α_1_ variant]). This haplotype is the same as what Bloom (2021) suggested to be important in rooting the SARS-CoV-2 phylogeny, which led us to consider five alternative scenarios based on *TopHap*, Kumar *et al*. (2021), Bloom (2021), Nextstrain, and PANGO (**Fig. 7**) involving eight positions that experienced mutations (α_1_-α_3_, β_1_-β_3_, ν_1_-ν_2_, and *x*) and gave rise to seven major early haplotypes. Our evaluation of these five scenarios is the most detailed comparison to date because of the size of the dataset analyzed and the variants included. For example, ν_1_ and ν_2_ variants were absent in the analysis by Bloom (2021), because genomes only sampled until the end of January 2020 were included, and variant *x* was missing from Kumar *et al*. (2021) analysis because its global frequency was less than 1% in the 68KG dataset.

**Figure 7.**
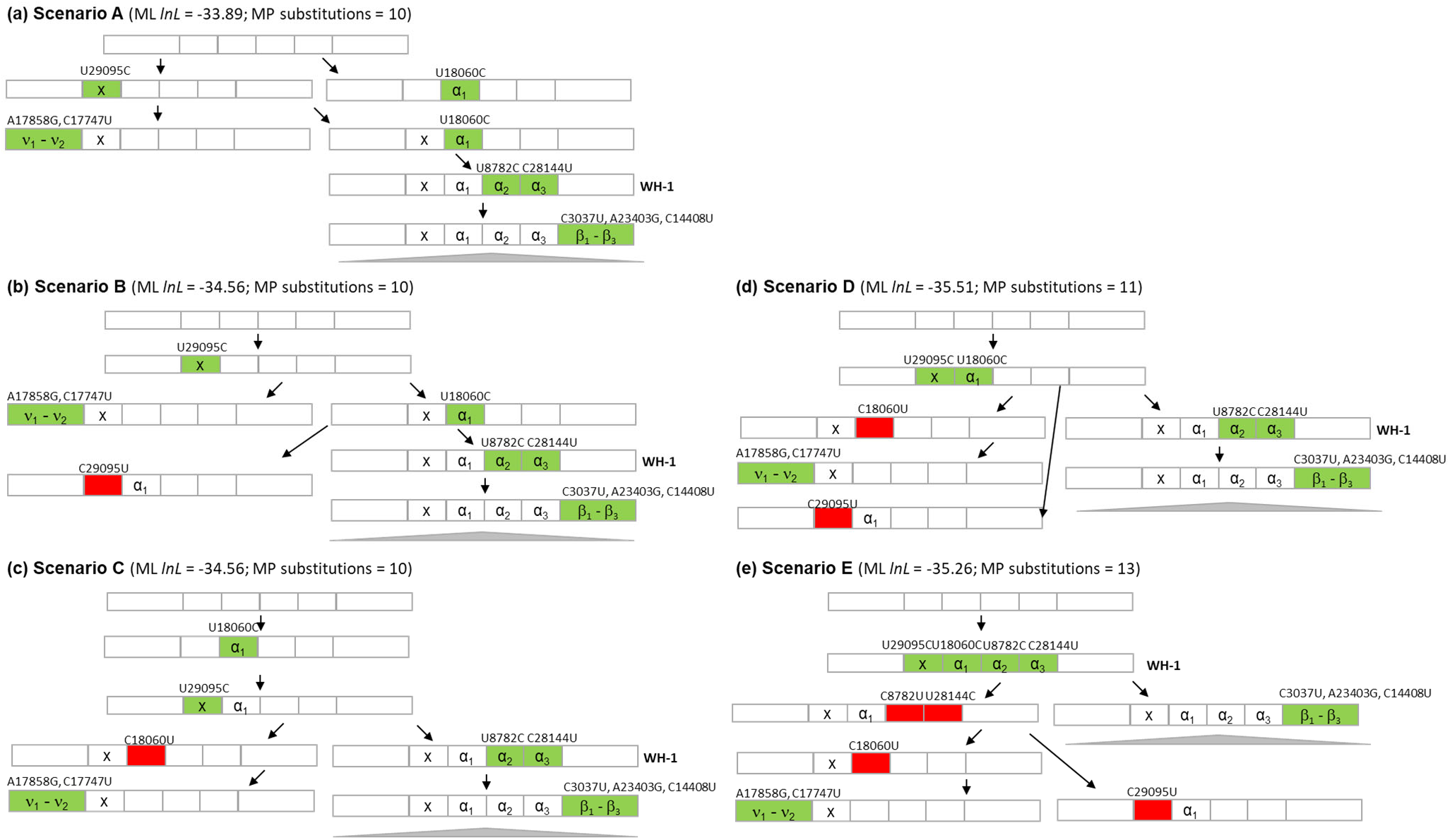
The early history of SARS-CoV-2 variants. Five root positions are explored in which the haplotype with mutation *x* has been added to the *TopHap* phylogeny in **figure 3** (A and B), Kumar *et al*., (2021) mutational history (B), Bloom, (2021) phylogeny (B and C), PANGO classification (D), and the Nextstrain classification (E). Haplotypes have eight positions that contain variants α_1_-α_3_. β_1_-β_3_. ν_1_-ν_2_,. and *x*. Genomic positions are shown whenever a mutation occurs: green box for forward and red for backward mutations). ML log likelihoods (*lnL*) and number of MP substitutions are shown. WH-1 is the haplotype corresponding to the Wuhan-1 genome. The gray triangle represents all the other SARS-CoV-2 haplotypes of the ongoing infections in the world.

We evaluated these five scenarios (topologies) using MP and ML optimally criteria (**Fig. 7**). We used the key seven haplotypes with the eight positions (α_1_-α_3_, β_1_-β_3_, ν_1_-ν_2_, and *x*). MP suggests scenarios A, B, and C to be equally parsimonious, and D and E (PANGO and Nextstrain, respectively) to be less parsimonious by 1 and 3 mutations. Scenario D and E were also less likely than A, B, and C, where we estimated the loglikelihood (*lnL*) of all five scenarios (topologies) using a GTR model of nucleotide substitutions in MEGA for the haplotypes shown in **Fig. 7**. While the log-likelihood of scenario A was the highest, it was only slightly higher than that for B and C that were equally likely. Among scenarios A, B, and C, *x* variant was lost in B, while the α_1_ variant was acquired twice in A and lost in C.

In all the three equally most parsimonious scenarios (A, B, and C), the addition of mutation *x* pushes back the MRCA of SARS-CoV-2 by one mutation compared to the proCoV2 sequence of Kumar *et al*. (2021). In these cases, the number of differences between Wuhan-1 and the MRCA is four (**Fig. 7**). With a mutation rate range of 6.64 × 10^-4^–9.27 × 10^-4^ substitutions per site per year (Pekar *et al*., 2021), we can estimate that proCoV2 existed 7.7–10.8 weeks before the December 24, 2019 sampling date of Wuhan-1. This places the progenitor of SARS-CoV-2 to have evolved in mid-September to Early-October 2019, which is earlier than the mid-November 2019 date favored by Pekar et al. (2021). Pekar et al. used rooting in scenario D in which the lineage containing (α_2_-α_3_ and β_1_-β_3_; PONGO B) is a sister group of the lineage containing α_1_ and ν_1_-ν_2_ (PONGO A). As noted above, this scenario receives lower bootstrap support than the alternative in which PONGO B arose from the ancestor containing α_1_. In this sense, Pekar et al. (2021) may have dated a later event in the SARS-CoV-2 phylogeny.

## 4 Conclusions

The ongoing global efforts to monitor the evolution of the SARS-CoV-2 coronavirus have motivated hundreds of laboratories worldwide to generate genome sequences continuously. The number of genomes has grown quickly, becoming orders of magnitude greater than the genome size. Rapid growth, low sequence variability, and the presence of sequencing error have made the direct use of phylogenetic methods on genome alignments challenging for such data (e.g., Morel *et al*., 2020). We have shown that the *TopHap* phylogeny for common variants and haplotypes in the 68KG SARS-CoV-2 dataset works well and agrees with the mutation tree produced using a mutation order approach (MOA) (Kumar *et al*., 2021). But, the *TopHap* approach offers some advantages over MOA. Firstly, MOA assumes the sequencing error rate to be constant throughout the outbreak, which is unlikely to hold for pathogenomic datasets acquired in different laboratories at different times. Secondly, MOA analysis needs to have mutant bases indicated at the outset, a limitation addressed by Kumar *et al*. (2021), but at a large computational expense. In contrast, *TopHap* analyses directly use outgroup in standard phylogenetic analysis. *TopHap* analysis is certainly more computationally efficient as the analysis of the 68KG dataset took only a few hours. In contrast, MOA took more than a week to compute.

Thirdly, *TopHap* analysis can use well-established methods to infer phylogeny and ancestral sequences to identify recurrent and backward mutations. In contrast, MOA assumes an infinite site model and, thus, is not suitable for detecting recurrent and backward mutations. Lastly, rarer haplotypes can also be attached to a backbone of a *TopHap* phylogeny by simply adding the genomic position of interest in constructing the MSA of haplotypes, as demonstrated above.

In conclusion, *TopHap* is a simple and effective method to build haplotype phylogenies and assess their statistical robustness. *TopHap* can be applied in any data containing a large number of sequences with a handful of variants, including other pathogens and tumor single-cell sequencing data that is now producing a large number of somatic cell sequences (Navin, 2015).

## Acknowledgments

We thank Sudip Sharma for comments and everyone depositing genome data on GISAID (list at http://igem.temple.edu/COVID-19).

## Author contributions

S.K. and S.M developed the original method, designed research; and implemented the technique; M.A.C.O., S.M., T.D., and Q.T. performed analyses; S.P. and S.W. assembled sequence alignments; and S.K., S.M., M.A.C.O., Q.T., and S.P. wrote the paper.

